# Growth differentiation factor-15 is associated with age-related monocyte immunosenescence

**DOI:** 10.1101/2020.02.05.935643

**Authors:** Brandt D. Pence, Johnathan R. Yarbro, Russell S. Emmons

## Abstract

**Background:** Immunosenescence is an age-associated decrease in function of immune cells precipitated by a variety of mechanisms and affecting nearly every immune cell subset. In myeloid cell subsets, aging reduces numbers of phagocytes and impairs their functional abilities, including antigen presentation, phagocytosis, and bacterial clearance. Recently, we have described an aging effect on several functions indicating immunosenescence in monocytes, including impaired mitochondrial function and reduced inflammatory cytokine gene expression during stimulation with lipopolysaccharide (LPS). We hypothesized that circulating factors altered by the aging process underly these changes. Growth/differentiation factor-15 (GDF-15) is a distant member of the transforming growth factor-beta superfamily that has known anti-inflammatory effects in macrophages and has recently been shown to be highly differentially expressed during aging. We used biobanked serum and plasma samples to assay circulating GDF-15 levels in subjects from our previous studies and examined correlations between GDF-15 levels and monocyte mitochondrial function and inflammatory responses.

**Results:** Monocyte interleukin-6 production due to lipopolysaccharide stimulation was negatively correlated to plasma GDF-15 levels. Additionally, serum GDF-15 was positively correlated to circulating CD16+ monocyte proportions and negatively correlated to monocyte mitochondrial respiratory capacity.

**Conclusions:** The results of these analyses suggest that GDF-15 is a potential circulating factor affecting a variety of monocyte functions and promoting monocyte immunosenescence, and thus may be an attractive candidate for therapeutic intervention to ameliorate this.

## INTRODUCTION

Aging is a multifactorial process leading to disruption of physiological function at subcellular, cellular, tissue, and organismal levels. As such, aging is the largest risk factor for most major chronic diseases [1]. Underlying these changes, at least in part, are the dual concepts of **inflammaging** and **immunosenescence**. Inflammaging is a generalized, age-associated, and chronic low-grade inflammatory state which is associated with disease progression and is included in the now well-established Hallmarks of Aging [2] and Pillars of Aging [3] concepts. Immunosenescence is a generalized decline in immune function which leads to increased susceptibility to infection and dysregulated responses to immunological stimuli [4]. Because inflammation is integral to immune responses, these processes have been suggested to share common etiology [4], but mechanisms linking the two conditions are largely unknown.

In the past several years, our laboratory has examined the effects of the aging process on immunosenescence of the innate immune system, with a focus on dysregulation of immunometabolism in monocytes [5–7]. We were the first to demonstrate impaired mitochondrial function in aged classical monocytes [5], and we have also found age-associated reductions in inflammatory cytokine responses to lipopolysaccharide (LPS) [6] and alterations in proportions of circulating monocyte subsets [5] that support previous and subsequent reports [8–12].

During the course of these studies, a large proteomics study was published which comprehensively profiled aging-related alterations in proteins in circulation [13]. Perhaps the most notable finding from this study was the high differential expression of growth differentiation factor-15 (GDF-15) between older and younger participants. Being generally unfamiliar with GDF-15, we performed a literature search and found that the protein, which is a member of the transforming growth factor-β superfamily, was originally described as macrophage inhibitory cytokine-1 [14] and was shown to be released from macrophages during inflammatory activation (thus potentially linking it to inflammaging) and to subsequently suppress inflammatory activation in macrophages (thus potentially linking it to innate immunosenescence). Moreover, GDF-15 is a constituent of the senescence-associated secretory phenotype (SASP) [15, 16], the proteome secreted by senescent cells [17] which is itself a major promoter of inflammaging [18].

Given these effects, we became interested in determining if GDF-15 is a link between cellular senescence, inflammaging, and innate immune system immunosenescence. During the course of our previous studies, we had biobanked blood plasma from older and younger adult participants, and therefore we assayed GDF-15 levels in these samples and conducted a small secondary analysis of our existing datasets to test the hypothesis that GDF-15 is correlated to indications of monocyte immunosenescence. Although the results of these analyses are associative, they support a potential link between GDF-15 and age-related monocyte dysfunction. Given the great interest in GDF-15 resulting from recent proteomics studies, and the paucity of data demonstrating age-associated biological effects of the protein, the outcomes reported here are an important preliminary step in the establishment of GDF-15 as a link between cellular senescence and immune system dysfunction resulting from the aging process.

## METHODS

### Subjects

Younger (18-35 yr) and older (60-80 yr) adults were recruited from the greater Memphis area for testing. Subject characteristics and inclusion/exclusion criteria have been previously described [6]. Subject characteristics are reported in **Table 1**, which was adapted from cohort 1 data from the corresponding table in our previous paper [6]. All subjects completed informed consent documents prior to enrollment, and all protocols were approved in advance by the Institutional Review Board at the University of

**Table 1:** Demographic and anthropometric characteristics of subjects.

|  | **Cohort 1** |  | **Prob** |
| --- | --- | --- | --- |
|  | **Aged (N=9)** | **Young (N=9)** |  |
| Age, yr (range) | 65.0 ± 1.2 (61-71) | 25.7 ± 1.9 (18-33) | p<0.001 |
| Height, cm (range) | 176.7 ± 3.0 (164-192) | 169.2 ± 4.1 (157-189) | p=0.166 |
| Weight, kg (range) | 79.6 ± 3.6 (57-93) | 77.3 ± 8.6 (48-129) | p=0.814 |
| BMI, kg/m<sup>2</sup> (range) | 25.5 ± 1.1 (20-32) | 26.4 ± 1.9 (19-36) | p=0.688 |
| Female, N (%) | 4 (44) | 5 (56) | p=0.637 |
| White, N (%) | 5 (56) | 6 (67) | Race: p=0.415 |
| Black, N (%) | 4 (44) | 2 (22) |  |
| Hispanic, N (%) | 0 (0) | 1 (11) |  |

Yr: year; cm: centimeters; kg, kilograms; BMI, body mass index; N, number of subjects.

Memphis (protocol #4361). Groups did not differ in any measured demographic or anthropometric characteristic, with the exception of age (p < 0.001), as designed.

### Monocyte functional assays

Detailed methods for functional assays for isolated human monocytes are reported in their respective papers [5, 6] and are briefly summarized here. Monocytes were isolated by immunomagnetic negative selection with CD16 depletion using a commercially-available kit, yielding an untouched population of classical (CD14+CD16-) monocytes at > 90% purity. Isolated monocytes were immediately used in downstream assays and were not frozen. For analysis of mitochondrial function, a Seahorse cell mito stress test assay was performed on 1.5×10^5^ monocytes per well in duplicate as described previously [5]. For determination of monocyte cytokine responses, isolated monocytes were stimulated for 24 hr with 1 ng·ml^-1^ LPS, then lysed with Trizol as previously described [6]. Isolated mRNA was purified using manufacturer’s instructions and reverse-transcribed to cDNA using a commercial kit, and cytokine fold expression was quantitated by qPCR using 4 ng cDNA assayed in duplicate against *B2M* as a housekeeping gene. For monocyte phenotype determination, 100 μl whole blood was stained with anti-CD14-PE and anti-CD16-Brilliant Violet 421 antibodies and analyzed on an Attune NxT flow cytometer as previously described [5]. Monocytes were gated based on forward- and side-scatter and partitioned into classical (CD14^+^CD16^-^), intermediate (CD14^+^CD16^+^), and non-classical (CD14^low^CD16^+^) subtypes based on cell surface expression of CD14 and CD16.

### GDF-15 analysis

For analysis of plasma cytokine levels, whole blood was collected into EDTA vacutainer tubes by venipuncture, and plasma was separated from erythrocytes by centrifugation at 1,500×*g* for 15 min at 4°C. Isolated plasma was aliquoted and frozen at −80°C until analysis. Circulating GDF-15 levels in plasma were determined by commercial ELISA (DY957, R&D Systems, Minneapolis, MN) according to manufacturer’s instructions. Plasma samples were diluted 10× in reagent diluent and assayed in duplicate against a standard curve. Intra-assay coefficient of variation was 7.7%. All samples were assayed on a single plate.

### Data analysis

All analyses were conducted in R v. 3.5.1 (R Foundation for Statistical Computing, Vienna, AUT). Demographic data with categorical outcomes (race, sex) were analyzed by Pearson’s chi-square test. Continuous demographic and anthropometric data, as well as monocyte functional data, were measured by independent samples T-test with one exception. Welch’s correction was applied to the T-test for *IL6* gene expression, as the data did not meet the criteria for homoscedasticity by Levene’s test. For determination of the associations between circulating GDF-15 levels and monocyte functional responses, bivariate Pearson’s correlations were performed. The significance level for all tests was set *a priori* at p ≤ 0.05. Reported results are mean ± SEM.

## RESULTS

All data used for analyses are available in a dedicated FigShare repository [19].

### Monocyte functional data

Monocyte functional data in **Figure 1** are adapted from our previous reports [5, 6] and demonstrate that monocytes from older adults have reduced mitochondrial respiratory capacity (**Figure 1A**), impaired cytokine responses to LPS (**Figure 1B**), and altered proportions of CD16^+^ (sum of intermediate and non-classical) subtypes (**Figure 1C**) compared to monocytes from younger adults. These data are reprinted here to aid in interpretation of subsequent correlational analyses.

**Figure 1.**
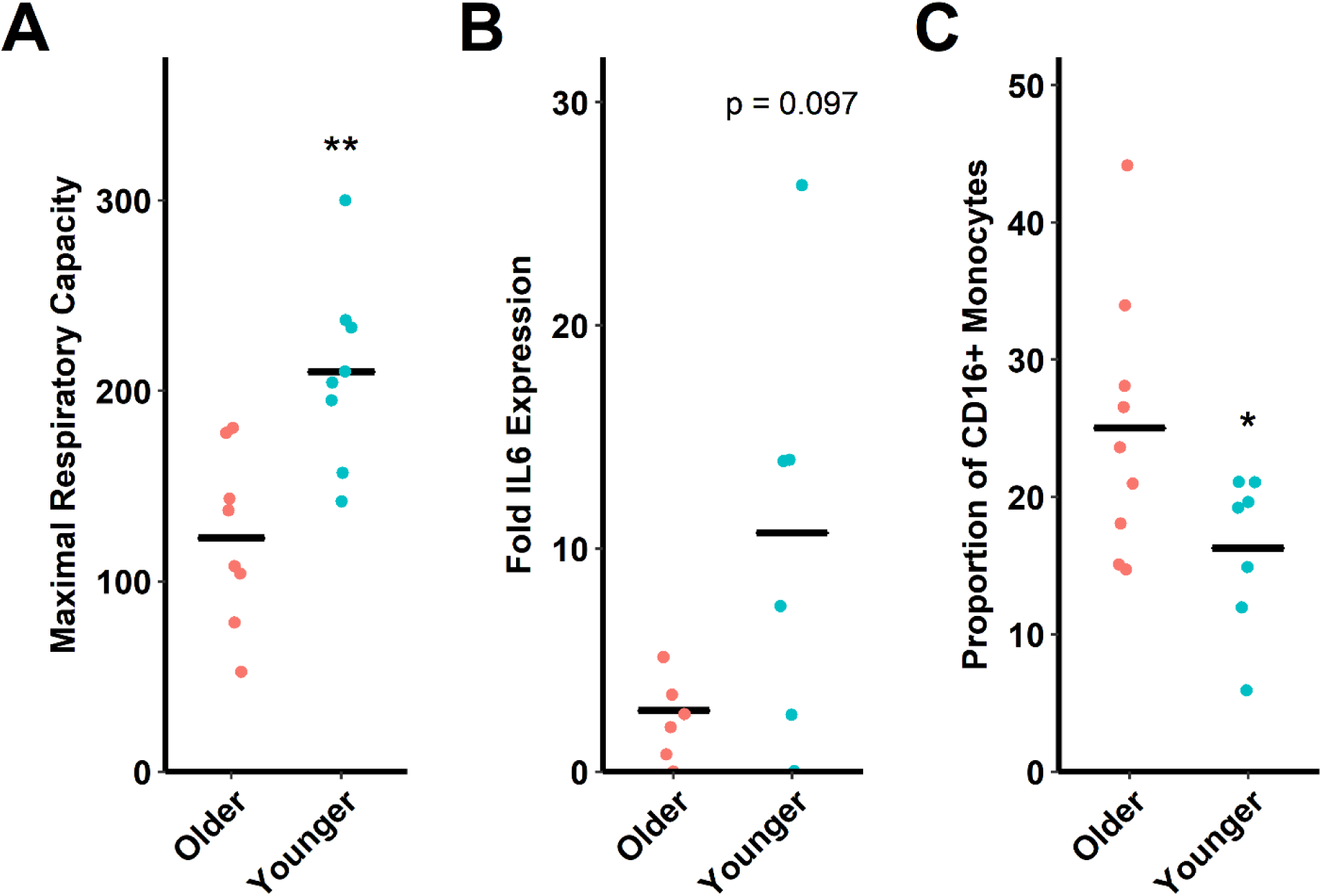
Indications of monocyte immunosenescence in older versus younger adults. (**A**) Older adults (60-80 yr) have reduced mitochondrial respiratory capacity compared to younger adults (18-35 yr). Maximal respiratory capacity is in pmol O_2_·min^-1^·(10^5^ monocytes)^-1^ as measured by a Seahorse XFp analyzer. (**B**) Older adults have a trend toward reduced *IL6* gene expression following LPS stimulation (by 2^-ΔΔCt^ method against *B2M*) compared to younger adults. (**C**) Older adults have increased circulating proportions of CD16+ monocytes (as percent total circulating monocytes) compared to younger adults. This subpopulation encompasses the sum total of proportions of intermediate and non-classical monocytes. * p ≤ 0.05 compared to older adults. ** p < 0.01 compared to older adults. N=6-9/group depending on assay.

### Plasma GDF-15

It was previously reported that GDF-15 is one of the most significantly upregulated circulating proteins during the aging process [13]. As expected, circulating GDF-15 levels in plasma from older adults was elevated nearly 3-fold in older compared to younger adults in the present study (t11 = 5.3321, p < 0.001, **Figure 2**).

**Figure 2.**
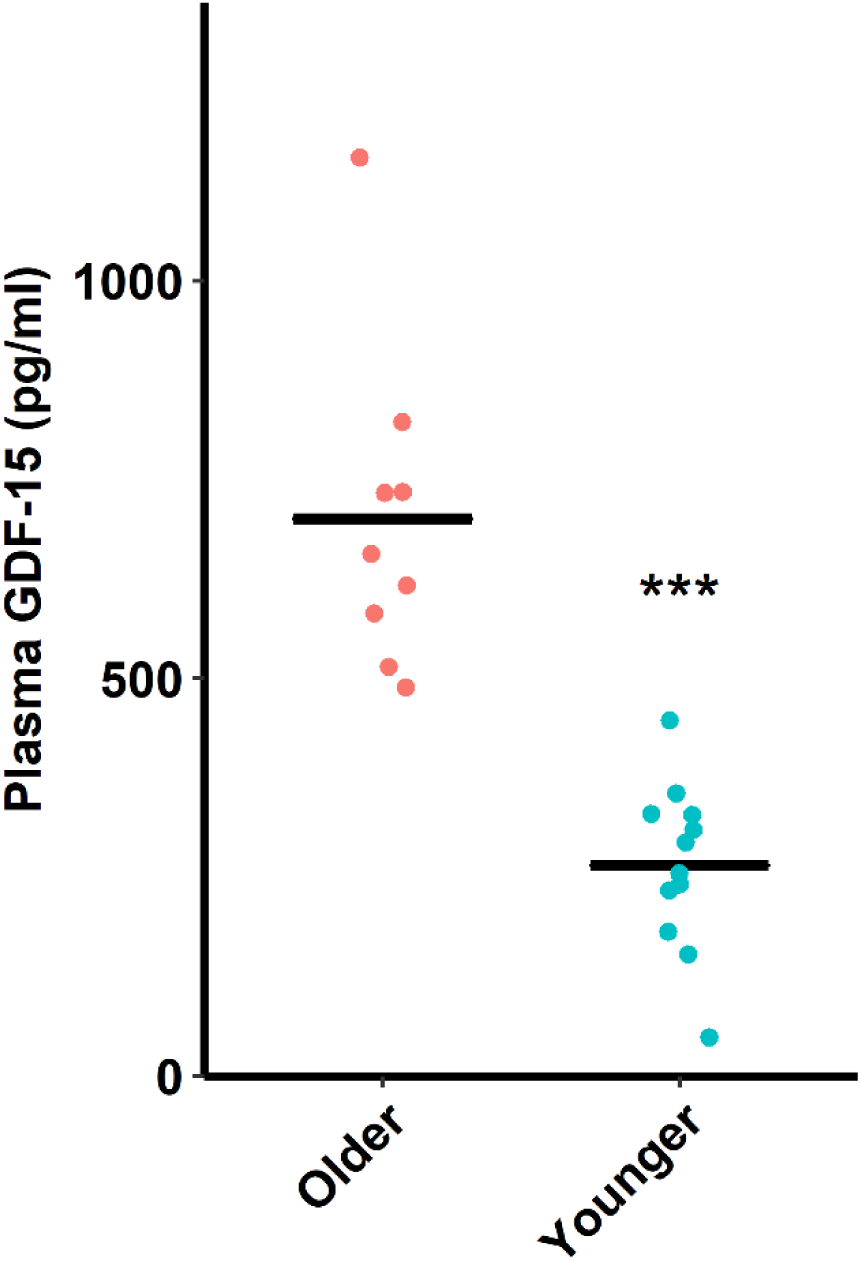
Older (60-80 yr) adults have increased plasma GDF-15 levels compared to younger (18-35 yr) adults. *** p < 0.001. N=9-12/group.

### Correlations

Given the established immunomodulatory properties of GDF-15 [14] and its upregulation with age, we reasoned that circulating levels of GDF-15 might be associated with immunosenescence. Our ability to conduct secondary analysis using our previous monocyte functional data, coupled with our biobanked human plasma samples, allowed us to conduct correlational analyses between circulating GDF-15 and indices of monocyte immunosenescence.

Circulating GDF-15 had strong negative correlation to monocyte maximal respiratory capacity (R = −0.747, p < 0.001, **Figure 3A**). GDF-15 had moderate negative correlation to *IL6* expression levels after LPS stimulation (R = −0.561, p = 0.046, **Figure 3B**), and had non-significant moderate negative correlation to *IL1B* and *IL10* expression (p > 0.05, not shown). Finally, GDF-15 had moderate positive correlation to circulating CD16+ monocyte proportion (sum of intermediate and non-classical monocyte proportions, R = 0.571, p = 0.021, **Figure 3C**) and moderate negative correlation to circulating classical (CD14+CD16-) monocyte proportion (R = −0.534, p = 0.033, not shown).

**Figure 3.**
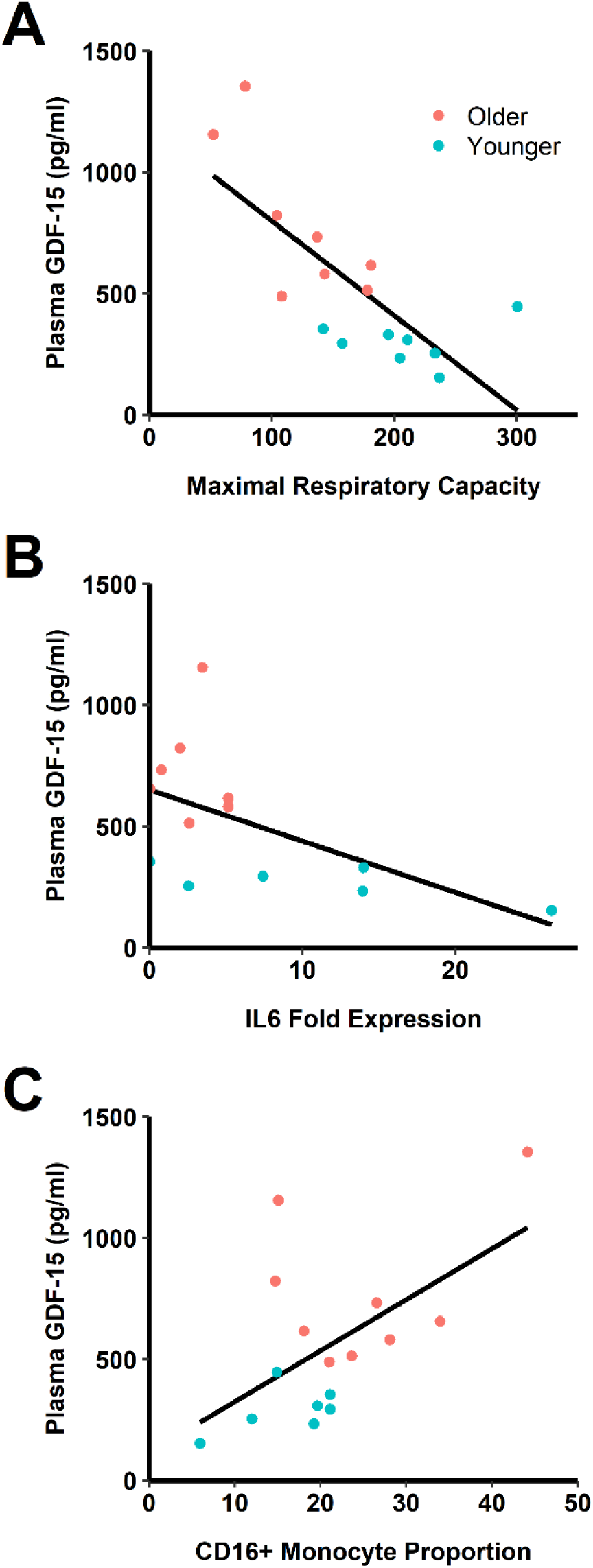
Associations between circulating GDF-15 and indices of monocyte immunosenescence. (**A**) GDF-15 has a strong negative correlation with maximal mitochondrial respiratory capacity (R = −0.747, p < 0.001). Maximal respiratory capacity is in pmol O_2_·min^-1^·(10^5^ monocytes)^-1^ as measured by a Seahorse XFp analyzer. (**B**) GDF-15 has a moderate negative correlation with *IL6* fold expression following LPS stimulation (by 2^-ΔΔCt^ method against *B2M*, R = −0.561, p = 0.046). (**C**) GDF-15 has moderate positive correlation with CD16+ monocyte proportions (R = 0.571, p = 0.021). N = 6-9/group depending on assay.

## DISCUSSION

Impaired immune responses are a normal consequence of aging, and increasing evidence suggests a key role of monocytes in these processes. Our lab has reported aging-induced mitochondrial dysfunction in monocytes, altered monocyte population distributions, and attenuated inflammatory responses following LPS stimulation [5–7]. A subset of these previously reported data were used for the analyses reported here.

GDF-15 is a cytokine member of the transforming growth factor-β superfamily and known to dampen or attenuate innate immune cell responses to a variety of stimuli. GDF-15 is released from human derived macrophages following TNF-α and IL-1β stimulation and in turn inhibits LPS stimulated TNF-α production in macrophages [14]. GDF-15 was recently observed to increase lipid accumulation and impair autophagy in oxLDL laden macrophages [20]. Furthermore, tumor-derived GDF-15 inhibits the synthesis of nitric oxide and TNF-α through TGF-activated kinase-1 in macrophages [21]. Thus, it appears GDF-15 blunts macrophage responses from diverse stimuli, but the common underlying mechanism by which GDF-15 attenuates innate immune cell response is still unknown.

Through secondary analyses of existing samples, we found upregulated circulating GDF-15 in older adults whose monocytes were tested for functional responses in our previous studies. GDF-15 has recently been identified as a core member of the senescence associated secretory phenotype [22] and has been shown to increase in peripheral circulation with age [13]. Serum GDF-15 was reported as a strong predictor of mortality in a Swedish male cohort [23], as well as of all-cause mortality in community-dwelling older adults [24]. The ∼3-fold increase in serum concentrations of GDF-15 in older subjects in our study is consistent with these previous reports demonstrating elevated secretions of GDF-15 with aging.

While the exact role of GDF-15 in the aging process is unclear, elevated GDF-15 has been observed in other chronic inflammatory disease including cardiovascular disease [25], obesity [26], type 2 diabetes [27], and cancer [28]. Together these data highlight a potential role for GDF-15 in chronic inflammatory diseases and mortality. These conditions are commonly associated with immune and mitochondrial dysfunction, but additional research is necessary to uncover the precise mechanisms by which GDF-15 participates in the pathogenesis of these diseases, and to establish whether or not it is elevated as a protective agent in pathology or is actively contributing to it.

In the present study we report, for the first time, associations between GDF-15 and indices of age-related immunosenescence. Using a secondary analysis of existing monocyte immunosenescence data paired with newly-acquired circulating GDF-15 protein data, we found significant negative correlations between GDF-15 levels and monocyte maximal respiratory capacity and LPS-stimulated IL-6 gene expression. We also demonstrated a significant positive correlation between GDF-15 and CD16^+^ monocytes proportions in peripheral circulation. While associative data must be interpreted cautiously, these data suggest a possible link between aging, GDF-15, and monocyte (dys)function.

Mitochondrial integrity and function are critical for monocyte and macrophage homeostasis. In this study, we observed a negative association between plasma GDF-15 levels and maximal mitochondrial respiratory capacity in monocytes. Respiratory capacity is the ability to maximally generate ATP through mitochondrial respiration [29]. The reduced respiratory capacity we demonstrated in our previous study [5] may render monocytes unable to generate sufficient ATP under aerobic conditions, leading to cellular dysfunction, although currently this is speculative. Because GDF-15 is thought to be a biomarker for mitochondrial dysfunction [30, 31] and indeed promotes mitochondrial function in macrophages [32], it is likely that GDF-15 is not causally-related to mitochondrial dysfunction in monocytes, but circulating levels of the protein may serve as a useful proxy measure for myeloid mitochondrial dysfunction.

Finally, non-classical CD16^+^ monocytes are typically characterized as anti-inflammatory and necessary for vascular maintenance [33]. However, non-classical monocytes have been observed to adopt a pro-inflammatory phenotype basally and to display features akin to cellular senescence in other cell types [34]. GDF-15 has been shown to promote cellular senescence in endothelial cells [35], and thus may play some role in the transition of monocytes to the non-classical phenotype which is increased with age [5, 8–11].

## Conclusions

Here we report associative data suggesting a potential role for GDF-15 in aging-induced monocyte dysfunction. Circulating GDF-15 levels were significantly correlated to impaired mitochondrial function, reduced LPS-stimulated inflammatory cytokine gene expression, and increased proportion of CD16+ circulating monocytes. While these correlational data are insufficient to demonstrate a direct effect of GDF-15 on monocyte immunosenescence, our findings highlight a potential role for this protein in regulating age-associated alterations in immune function.

## DECLARATIONS

### Ethics Approval and Consent to Participate

All study activities were approved by the Institutional Review Board at the University of Memphis (protocol #4361). Subjects provided informed consent and were free to withdraw at any time.

### Consent for Publication

Not applicable

### Availability of Data and Materials

The datasets and analytical scripts generated and analyzed during the current study are available in the FigShare repository, https://doi.org/10.6084/m9.figshare.c.4844223.

### Funding

The study was supported by an American Heart Association grant (18AIREA33960189) to BP.

### Authors’ Contributions

BP conceived and designed the study. BP and JY collected the data. BP analyzed the data. BP and RE drafted the manuscript. BP, RE, and JY revised the manuscript drafts. All authors read and approved the final manuscript.

## Acknowledgements

The authors would like to acknowledge the participants in this study.

